# EGFRvIII-Targeted Virus-Like Particles Enable Selective Genome Editing and Elimination of Glioblastoma Cells

**DOI:** 10.1101/2025.08.26.672521

**Authors:** Canan Bayraktar-Odabas, Ibrahim Sahin Ozyildiz, Tugba Bagci-Onder

**Author notes:** Corresponding author Koç University School of Medicine, Rumelifeneri Yolu, 34450, Sarıyer, Istanbul, Türkiye.

## Abstract

Glioblastoma (GBM) is a highly aggressive malignancy with poor prognosis, frequently driven by aberrant signaling through the mutant epidermal growth factor receptor variant III (EGFRvIII). This unique tumor-specific alteration provides an attractive opportunity for precision therapies that can discriminate malignant from normal tissue. In this study, we developed a modular virus-like particle (VLP) platform engineered to selectively recognize EGFRvIII-positive cells while minimizing off-target activity. By systematically screening single-chain variable fragments (scFvs) and peptide ligands displayed on engineered viral envelopes, we identified an optimal targeting configuration that maximized specificity without compromising entry efficiency. Beyond targeting, we optimized multiple layers of VLP design—including packaging stoichiometry and gRNA backbones—to achieve robust encapsidation and delivery of Cas9 ribonucleoproteins (Cas9-RNPs). Functional assays demonstrated efficient genome editing in reporter systems and confirmed the capacity of our platform for reliable therapeutic cargo delivery. Most importantly, EGFRvIII-targeted VLPs translated delivery into therapeutic outcomes, enabling potent and highly selective elimination of EGFRvIII-positive glioblastoma cells while sparing non-target cells. Collectively, this work establishes a versatile and programmable framework for tumor-targeted VLP therapeutics and lays the foundation for future in vivo studies toward precision treatment of glioblastoma and other EGFRvIII-driven cancers.

## INTRODUCTION

Glioblastoma (GBM) is the most aggressive primary brain tumor in adults, characterized by extensive heterogeneity, diffuse infiltration, and resistance to conventional therapies [1, 2]. Despite maximal surgical resection followed by radiotherapy and temozolomide chemotherapy, median survival remains only 14–18 months [3]. The limited efficacy of standard regimens has motivated the search for precision strategies that selectively eliminate tumor cells, while sparing normal brain tissue. One of the most promising tumor-specific alterations in GBM is the epidermal growth factor receptor variant III (EGFRvIII), a deletion mutant expressed in ∼20–30% of patients and absent from normal tissues [4, 5]. Its prevalence and tumor-restricted expression have made EGFRvIII a major focus of immunotherapy and targeted delivery approaches [6, 7].

Virus-like particles (VLPs) and membrane-derived enveloped delivery vehicles (EDVs) have recently emerged as compelling solutions that combine the efficient cell entry of viral systems with the genome-free, transient release of nonviral methods [8-11]. In these systems, Cas9 ribonucleoproteins (RNPs) or other genome editors are pre-assembled and packaged into particles for “hit-and-run” activity, enabling efficient modification of primary and post-mitotic cells without stable vector integration [12-14]. A central determinant of both selectivity and efficiency is the envelope engineering strategy. Classical VSV-G pseudotyping confers broad tropism primarily through binding to the low-density lipoprotein receptor (LDLR), but it is now clear that VSV-G can also exploit other members of the LDLR family and even distinct receptors such as Lgr4. Notably, cells lacking LDLR remain susceptible to VSV-G–pseudotyped viruses, underscoring the multifactorial nature of VSV-G–mediated entry [15-17]. For therapeutic applications, entry must be restricted to disease-relevant targets. Accordingly, modular systems now incorporate antibody fragments (scFvs) or ligands onto envelope scaffolds, or redesign fusogenic proteins to decouple entry from LDLR, thereby achieving programmable, cell-type specificity [10, 18, 19].

Recent advances highlight the therapeutic potential of such targeting approaches. Doudna and colleagues demonstrated that EDVs displaying scFvs can program receptor-specific uptake and achieve *in vivo* human T cell engineering with Cas9-RNPs [10, 11]. Liu and colleagues established engineered virus-like particles (eVLPs) for efficient genome editing and further modularized envelope design for tailored delivery [8, 9]. Complementary approaches such as Nanoblades have achieved effective Cas9-RNP delivery into primary cells and tissues using retroviral scaffolds [13]. In parallel, antibody engineering has yielded scFvs with refined affinity and specificity, enabling more precise targeting of tumor-associated receptors including EGFRvIII [20, 21].

Leveraging these advances, we sought to develop a tumor-selective VLP platform for glioblastoma that couples EGFRvIII-directed uptake with CRISPR-Cas9 RNP delivery. Specifically, we (i) benchmarked two transmembrane scaffolds (ACE2-TM and VSV-TM) decorated with EGFRvIII-targeting scFvs or peptides to optimize entry efficiency and specificity; (ii) implemented a Gag–Cas9 fusion–based packaging scheme inspired by prior eVLP and Nanoblade systems to maximize RNP incorporation [8, 13]; and (iii) compared gRNA packaging plasmids, including a SuperBlade5 design, to enhance editing output per particle. Through these optimizations, we demonstrate that Cas9-RNP VLPs can achieve selective elimination of EGFRvIII(+) glioblastoma cells by targeting essential genes—establishing a modular and transient genome editing platform for tumor-specific intervention.

## MATERIALS and METHODS

### Plasmid Construction

Targeting modules, including three single-chain variable fragments; scFv1 [22], scFv2 [23], scFv3 [24] and peptide ligands; Peptide1 [25], Peptide2 [26], Peptide3 [27] (Supplementary table 1), were cloned into envelope scaffolds. For surface display, two different transmembrane scaffolds were used: ScFv-V, in which scFvs or peptides were fused to the typical VSV-G transmembrane/cytoplasmic tail domain, and ScFv-A, in which scFvs or peptides were fused to the ACE2-derived transmembrane/cytoplasmic tail that we previously showed to support efficient targeting in engineered VLPs[28]. scFv and transmembrane fragments were synthesized as DNA fragments (Twist Biosciences, USA), while short peptide sequences were synthesized as oligonucleotides Macrogen Inc. (Seoul, South Korea), phosphorylated using T4 polynucleotide kinase (NEB), and annealed to generate double-stranded inserts. All constructs were PCR-amplified using Phusion^®^ High-Fidelity DNA Polymerase (Thermo Fisher Scientific, Waltham, MA, USA). These fragments were cloned into the pcDNA3-sACE2(WT)-8his vector (Addgene #149268), replacing the HA-tag region from VSV-Gmut plasmid (Addgene #207317), while retaining the ACE2 signal and transmembrane domains. Ligation was conducted using T4 DNA ligase (NEB) following the manufacturer’s instructions, and constructs were transformed into *E. coli* Stbl3 competent cells. Successful clones were confirmed by Sanger sequencing.

### Cell Culture

Human Embryonic Kidney 293T cells, were purchased from American Type Culture Collection (ATCC) and EGFRvIII-positive DK-MG glioblastoma cells were kindly gifted by Dr. Malte Kriegs [29]. All cells were maintained in Dulbecco’s Modified Eagle Medium (DMEM; Gibco, USA) supplemented with 10% fetal bovine serum (FBS; Gibco, USA) and 1% penicillin–streptomycin (Gibco, USA) at 37 ^°^C in a humidified incubator with 5% CO_2_. Cells were routinely passaged when they reached approximately 70% confluency. DK-MG^+^ cells were used at low passage numbers, as they are reported to lose EGFRvIII expression upon repeated passaging. All cell lines were routinely screened for mycoplasma infection.

### Cell Line Generation and Validation

#### hROSA26 knock-in line

To generate targeted knock-in HEK293T cell lines, a donor plasmid containing a CAG-EGFP cassette flanked by homology arms to the human ROSA26 locus was co-transfected with two PX459-based plasmids encoding guide RNAs (targeting the top and bottom strands of the ROSA26 locus). Briefly, 2.5 × 10^6^ HEK293T cells were seeded in 10-cm plates one day before transfection. For each plate, 1.25 µg PX459-guide1, 1.25 µg PX459-guide2, and 2.5 µg donor plasmid were mixed with PEI (as described in the VLP production section). 48h post-transfection, cells were subjected to hygromycin selection (200 µg/mL). Surviving colonies were expanded and seeded into 96-well plates by limiting dilution to isolate single-cell– derived clones. Colonies exhibiting uniform EGFP expression under fluorescence microscopy (constant gain and exposure) were expanded for further validation.

#### Genotyping and validation

Genomic DNA was extracted, and locus specific PCR was performed using primers binding outside the homology arms. PCR cycling conditions were: 98 °C for 1 min; 34 cycles of 98 °C for 30 s, 67 °C for 30 s, 72 °C for 1 min; and a final extension at 72 °C for 4 min. Short extension times allowed distinction of heterozygous versus homozygous integration events. To confirm 5′ integration, an additional PCR was performed using a forward primer outside the 5′ homology arm and a reverse primer within the hygromycin cassette, using similar cycling conditions with an annealing temperature of 66 °C.

#### EGFRvIII-expressing line

To establish an EGFRvIII-expressing HEK293T line, the EGFP cassette of plasmid #20737 (Addgene) was replaced with a blasticidin resistance gene, generating a transfer plasmid for viral production. HEK293T cells were transduced with the resulting virus, and selection was applied with blasticidin (10 µg/mL) at 48h post-infection. EGFRvIII expression in the resistant population was validated by qPCR and Western blot analysis.

### VLP Production

VLPs were produced in HEK293T cells by transient transfection using polyethyleneimine (PEI; Polysciences, Cat#23966-1) at a DNA:PEI ratio of 1:4 (w/v). Cells were seeded one day prior to transfection at densities of 4 × 10^6^ cells per 10-cm dish or 1 × 10^6^ cells per well of a 6-well plate and transfected at ∼90% confluence. For standard VLP production, a total of 7.5 µg DNA was used per 10-cm dish (2.5 µg per well in 6-well format). Plasmid mixtures consisted of: (i) gRNA-backbone plasmid (10 parts), (ii) packaging plasmid (9 parts, psPAX2), and (iii) envelope plasmid (1 part). For scFv-or peptide-decorated VLPs, the DNA mixture included 1.25 µg pLEX-307-GFP, 1.125 µg psPAX2, 37.5 ng mutant VSV-G (Addgene, #207323), and 87.5 ng scFv1 or peptide- envelope plasmid per well. To determine optimal targeting stoichiometry, the ratio of mutant VSV-G to scFv1-V envelope was systematically varied from 100:0 to 0:100.

For Cas9–RNP loading, the Gag–Cas9 fusion system (Addgene #181752) was co-transfected with the packaging plasmid pUMVC (Addgene #8449), with DNA amounts adjusted according to the indicated ratios.

gRNA delivery was benchmarked using gRNA-Backbone1 [13, 30] (SuperBlade5; Addgene #134913), with later comparisons to gRNA-Backbone2, constructed in-house by cloning the U6 promoter and gRNA scaffold from lentiCRISPR v2 (Addgene #52961) into pUC19. Conventional lentiviral controls were produced using lentiCRISPR v2.

DNA–PEI mixtures were incubated for 20 min at room temperature before addition to cells. After 16 h, the transfection medium was replaced with fresh DMEM with FBS. Conditioned media were harvested at 48h and 72h post-transfection, clarified by centrifugation, and filtered through 0.45 µm syringe filters. To ensure that all envelope-coated VLPs were transduced at the same multiplicity of infection (MOI), particle titers were quantified using the QuickTiter™ Lentivirus Titer Kit (Cell Biolabs, San Diego, CA, USA) according to the manufacturer’s instructions, and MOI was calculated based on particle counts.

### Flow Cytometry

Flow cytometry was performed on a CytoFLEX Flow Cytometer (Beckman Coulter, Brea, CA, USA). Cells seeded in 24-well plates were washed with PBS and detached by incubation with 500 µL of PBS–EDTA (1%) per well for 3–5 min at 37 °C. Harvested cells were collected into 1.5-mL tubes, analyzed for GFP expression using the FITC channel. A minimum of 10,000 events per sample were recorded. Data acquisition was performed using the CytExpert software (Beckman Coulter), and analysis was carried out with the same platform. All measurements were performed in technical triplicates.

### Quantitative PCR (qPCR)

Total RNA was isolated using the NucleoSpin RNA Isolation Kit (Macherey-Nagel, Germany), and 500 ng was reverse-transcribed with the iScript^™^ cDNA Synthesis Kit (Bio-Rad, USA). qPCR was carried out with SYBR Green Master Mix (Roche, Switzerland) on a LightCycler^®^ 480 Instrument II (Roche, Switzerland). Relative gene expression was calculated using the ΔΔCt method, with GAPDH as the reference gene. Primer sets specific for EGFRvIII, GFP, and GAPDH were used.

### Western Blotting

For immunoblotting, VLP preparations were mixed with 4× Laemmli sample buffer (Bio-Rad, USA) supplemented with 2-mercaptoethanol (9:1 ratio) and heated at 95 °C for 10 min. Equal amounts of protein were resolved on Mini-PROTEAN^®^ TGX^™^ gradient precast gels (Bio-Rad, USA) and protein extracts were separated by electrophoresis at 120 V for 75 min. Proteins were transferred to PVDF membranes using the Trans-Blot^®^ Turbo^™^ system (Bio-Rad, USA). Membranes were blocked in TBS-T (0.1% Tween-20) containing 5% non-fat dry milk for 1 h at room temperature and then incubated overnight at 4 °C with primary antibodies diluted in TBS- T supplemented with 2% BSA and 0.02% NaN_3_. The following antibodies were used: anti-Cas9 (#7A9-3A3, Cell Signaling 1:1000), anti-VSV-G (#V5507, Sigma-Aldrich; 1:1000), anti-EGFR (#ab131498, abcam,1:1000), and anti-GAPDH (#ab9485, Abcam; 0.7–1 μg/mL). After three washes with TBS-T (5, 10, and 15 min), membranes were incubated for 1 h at room temperature with HRP-conjugated secondary antibodies (1:5000 dilution). Signal detection was performed using Pierce^™^ ECL Western Blotting Substrate (Thermo Fisher Scientific, USA), and chemiluminescence was visualized on an Odyssey^®^ Fc Imaging System (LI-COR Biosciences, USA).

### Cell Viability Assay

Equal numbers of EGFRvIII (+) and EGFRvIII (-) cells were seeded into black-framed, clear-bottom 96-well plates. The following day, cells were infected with VLPs carrying RPL9-targeting Cas9–RNPs, either coated with EGFRvIII-specific scFv1-A, scFv1-V or with wild-type VSV-G. As a control, cells were treated with non-targeting VLPs, which carried Cas9-RNPs with nontargeting gRNA. 16h after infection, the culture medium was replaced with fresh medium. Cell viability was measured at 3-, 5-, and 7-days post-infection using the MTT assay. At each time point, 25 µL of MTT solution (3 mg/mL in PBS; Sigma-Aldrich) was added to each well and incubated for 2 h at 37 °C. The medium was then removed, 100 µL of DMSO was added to solubilize the crystals, and absorbance was recorded at 570 nm using a microplate reader. Each condition was performed in technical triplicates.

### Statistical Analysis

Statistical analyses were performed using GraphPad Prism v9 (GraphPad Software, USA) and ImageJ (NIH, USA). Data are presented as mean ± standard deviation (SD) unless otherwise stated. Comparisons involving multiple parameters were evaluated by two-way analysis of variance (ANOVA). A two-tailed p-value < 0.05 was considered statistically significant. Specific details of statistical tests applied to each dataset are provided in the corresponding figure legends.

## RESULTS

### Design and optimization of EGFRvIII-targeted VLPs displaying scFvs or peptide ligands

To establish a platform for selective EGFRvIII targeting, we designed VLPs in which single-chain variable fragments (scFvs) or peptide ligands were fused to viral transmembrane domains (VSV-TM (scfv-V) or ACE2- TM (scFv-A)) for surface display (**Fig. 1A)**. To monitor entry efficiency in the initial proof-of-concept assays, VLPs were loaded with GFP-encoding RNA, allowing intracellular GFP signal to serve as a quantitative readout of uptake by flow cytometry. qPCR confirmed EGFRvIII expression in engineered HEK-293T cells, while the DK- MG glioblastoma line, which endogenously expresses EGFRvIII [29], served as a biologically relevant model **(Fig. 1B).** Western blot analysis further validated EGFRvIII protein expression in both cell types **(Fig. 1C).** We next evaluated targeting efficiency using three different scFvs and three peptide ligands displayed via either VSV-TM or ACE2-TM domains in both HEK-293T and DK-MG cells. Flow cytometry revealed that scFv1 consistently provided the strongest specificity between EGFRvIII (+) and EGFRvIII (−) populations. VSV-TM– based display yielded the highest overall entry efficiency, whereas ACE2-TM–based display enhanced specificity by reducing background uptake. In particular, ACE2-TM–scFv1 VLPs achieved at least a four-fold increase in uptake in EGFRvIII (+) cells compared to EGFRvIII (−) counterparts across both cell lines **(Fig. 1D).** To further refine the system, we optimized the plasmid ratios between mutant VSV-G and scFv constructs during VLP production. This titration revealed that a 70:30 ratio provided the most favorable balance between efficiency and specificity, strengthening the selective advantage of scFv1-coated particles **(Fig. 1E).** Together, these results demonstrate that scFv1-coated VLPs, particularly when displayed through ACE2-TM, enable efficient and selective entry into EGFRvIII-positive glioblastoma cells, providing a robust platform for targeted delivery.

**Figure 1.**
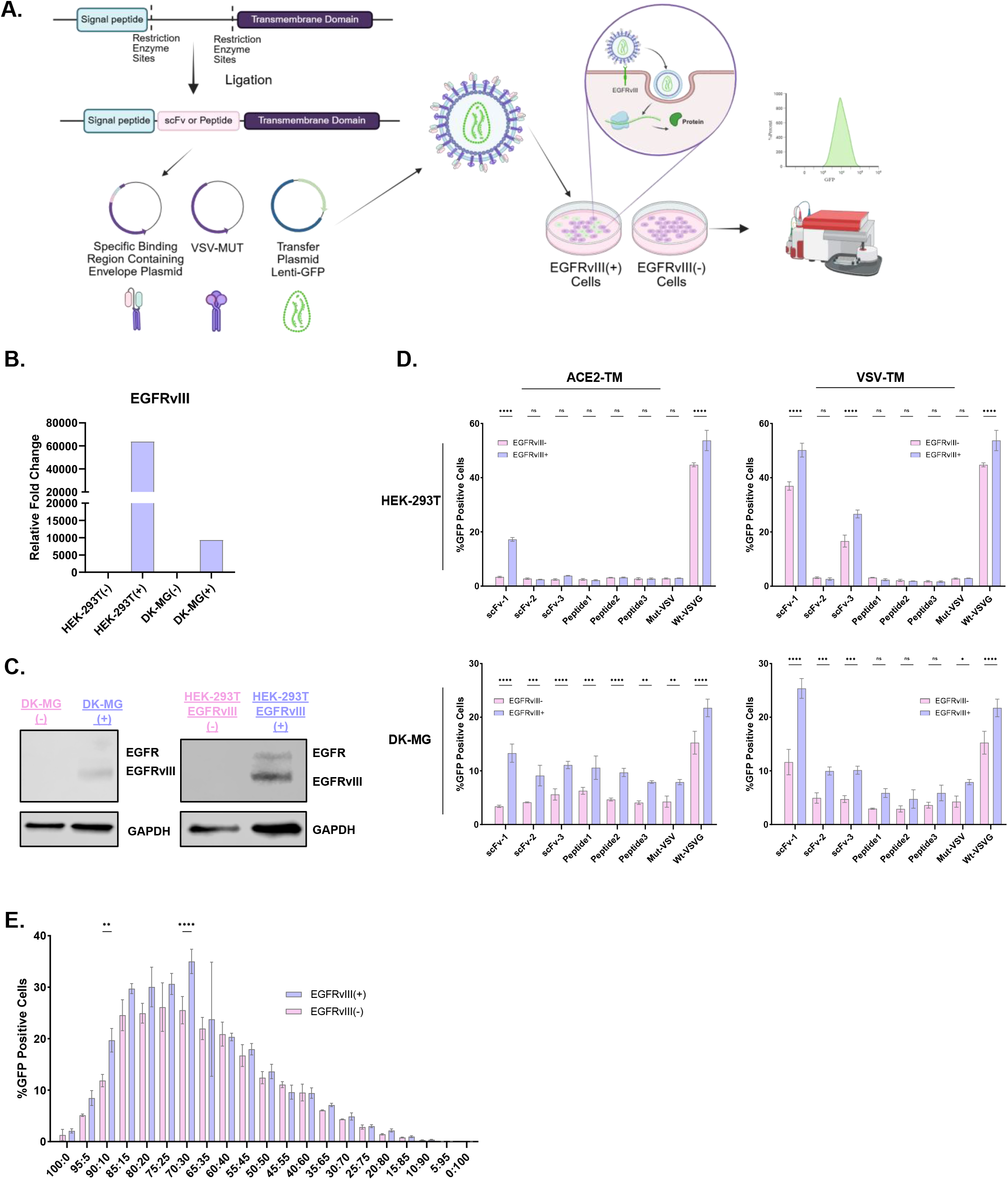
Design and validation of EGFRvIII-targeted VLPs displaying scFvs or peptide ligands via viral transmembrane domains. **A.** Schematic overview of the VLP design in which single-chain variable fragments (scFv) or targeting peptides are fused to viral transmembrane domains (VSV-TM or ACE2-TM) to enable their surface display. Engineered VLPs were applied to EGFRvIII(+) and EGFRvIII(−) cells, with GFP signal intensity measured by flow cytometry as a readout of entry efficiency. **B.** Relative EGFRvIII mRNA expression in engineered HEK-293T and DK- MG cell lines, as determined by qPCR. **C.** Western blot analysis confirming EGFR and EGFRvIII protein expression in DK-MG and HEK-293T cells. GAPDH was used as a loading control. **D.** Comparative evaluation of three different scFvs and three targeting peptides displayed via ACE2-TM or VSV-TM domains. Entry efficiency was assessed in both HEK- 293T and DK-MG cells using VLPs pseudotyped with wild-type VSV-G (WT-VSVG), mutant VSV-G (Mut-VSVG), or the designed targeting modules. **E.** Optimization of plasmid ratio between Mut-VSVG and scFv constructs during VLP production. Entry efficiency was quantified in HEK-293T EGFRvIII-positive and -negative cells. Statistical analysis was performed using two-way ANOVA. ^*^ p < 0.05, ^**^ p < 0.01, ^***^ p < 0.001, ^**^ p < 0.0001; ns: non-significant

### Optimization of Cas9-RNP packaging and delivery using GFP knock-in reporter cells

As a step toward developing VLPs capable of selective cell killing, we first optimized Cas9-RNP packaging efficiency. We employed a Gag–Cas9 fusion–based strategy to load Cas9-RNPs into VLPs **(Fig. 2A).** To quantify outcomes, we first generated and validated a homozygous GFP knock-in reporter line at the hROSA26 locus; where loss of GFP directly reports genome editing activity. Correct integration was confirmed by fluorescence microscopy and locus-specific PCR **(Fig. 2B–C).** This reporter system provided a robust and scalable readout, linking packaging design directly to functional editing outcomes.

**Figure 2.**
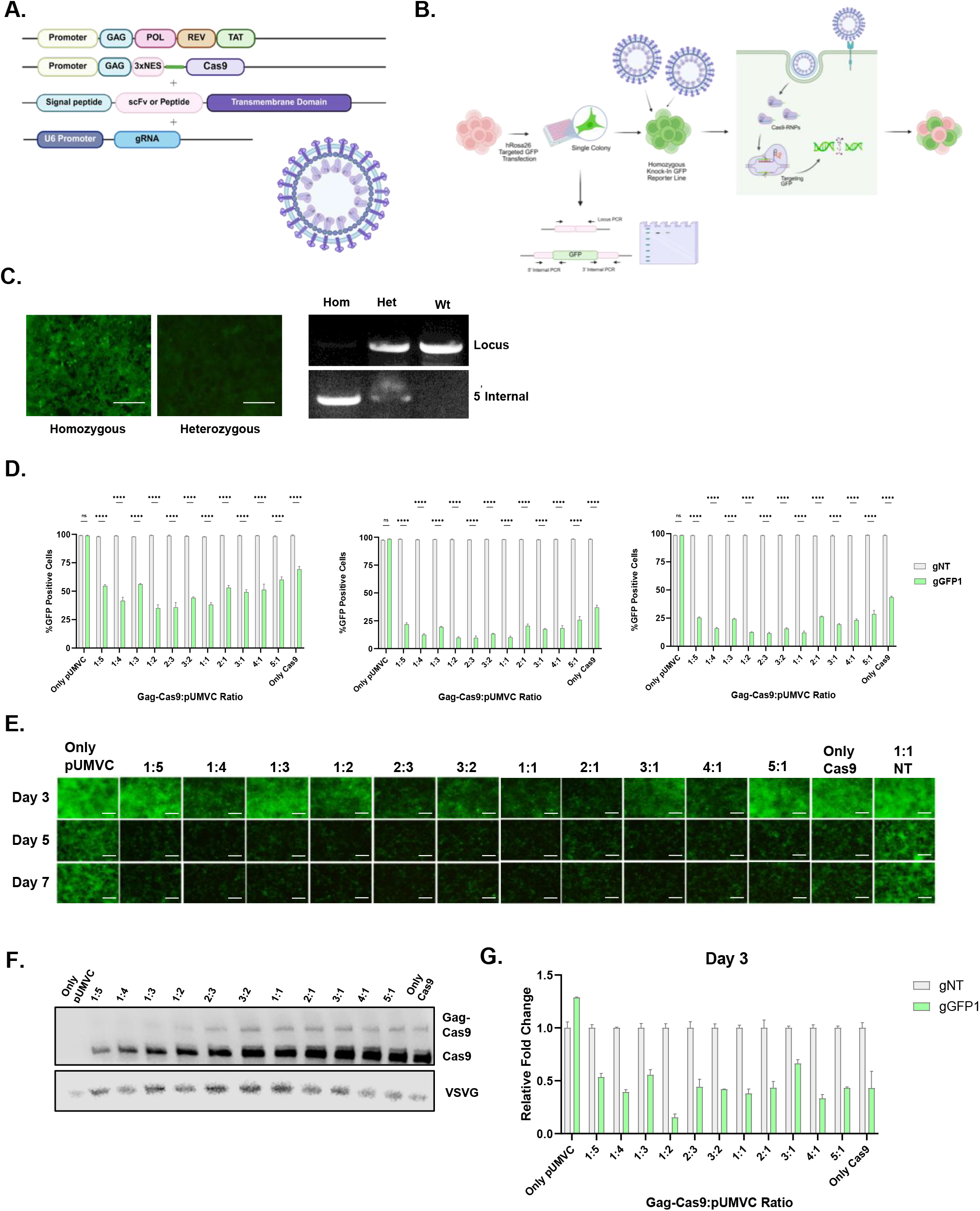
Optimization of Cas9-RNP packaging into VLPs for targeted genome editing. **A.** Plasmid constructs used for packaging Cas9-RNP into VLPs, including Gag, Cas9, and targeting modules. **B.** Experimental workflow using homozygous GFP knock-in reporter cells to assess genome editing following VLP delivery. **C.** Validation of GFP knock-in reporter cells by fluorescence microscopy and locus-specific PCR. **D.** Quantitative analysis of editing efficiency across 13 different Gag-Cas9:pUMVC plasmid ratios on days 3, 5, and 7, shown as bar plots. **E.** Representative fluorescence microscopy images corresponding to selected conditions from panel D, demonstrating relative GFP signal loss (scale bar: 200 µm). **F.** Western blot analysis of VLPs produced under the 13 ratio conditions, probing for Cas9 and VSV-G. **G.** qPCR-based quantification of editing efficiency for the 13 plasmid ratios at day 3. Statistical analysis was performed using two-way ANOVA. ^*^ p < 0.05, ^**^ p < 0.01, ^***^ p < 0.001, ^**^ p < 0.0001; ns: non-significant

Using this system, we systematically tested 13 different Gag-Cas9:pUMVC ratio combinations to modulate Cas9 stoichiometry during packaging. Cas9-RNP VLPs induced strong GFP loss across nearly all conditions **(Fig. 2D–E).** Notably, increasing Cas9 plasmid input did not yield proportional gains; instead, balanced ratios such as 1:1, 1:2, and 2:3 produced the most effective and reproducible editing. The imaging data clearly illustrate this effect, with robust and uniform GFP loss under balanced conditions but weaker and less consistent signal reduction at Cas9-rich ratios. Western blotting revealed a modest rise in Gag–Cas9 and free Cas9 signal with higher Cas9 input, but this was accompanied by comparatively lower VSV-G levels at Cas9-rich ratios **(Fig. 2F),** consistent with a stoichiometric imbalance that compromises functional entry rather than enhancing cargo availability. qPCR at day 3 independently validated the flow-based editing readouts across all ratios **(Fig. 2G),** confirming the reproducibility of these effects across assays. Our systematic survey across 13 conditions revealed that editing plateaus beyond balanced stoichiometry, with excess Cas9 compromising VLP integrity. This establishes a generalizable principle that rational tuning of cargo-to-structural protein ratios is essential not only for maximizing editing efficiency but also for ensuring particle quality, a consideration critical for therapeutic translation. Based on these observations, we adopted the 1:1 ratio for subsequent experiments, as it provided both robust editing and reliable particle composition.

### Evaluation of gRNA delivery backbones reveals an optimized design for VLP-mediated editing

To benchmark gRNA delivery strategies, we compared VLPs generated with two different gRNA backbones, gRNA-Backbone1 and gRNA-Backbone2, against a conventional lentiviral system (lentiCRISPR v2), in which both Cas9 and gRNA are delivered in the form of RNA **(Fig. 3A).** Flow cytometry revealed that Backbone1 consistently outperformed Backbone2 across all tested conditions, including both VLP input volumes (150 µl and 300 µl) and at all three time points (day 3, 5, and 7) **(Fig. 3B).** Under these input conditions, Backbone1- based VLPs also achieved higher editing efficiencies than lentiCRISPR v2. Representative fluorescence images confirmed these trends, showing a stronger reduction of GFP signal in Backbone1-treated cells compared with Backbone2 or lentiCRISPR v2 **(Fig. 3C).** qPCR quantification of GFP transcript levels was fully consistent with the flow cytometry data, further validating Backbone1 as the most effective design among the tested systems **(Fig. 3D).** Importantly, this comparison highlights that not all gRNA backbones are equally suited for RNP-based delivery. While lentiviral vectors traditionally rely on RNA-packaging cassettes, VLPs must efficiently load pre- formed Cas9–RNP complexes, placing different demands on gRNA stability and incorporation. Systematically benchmarking alternative backbones therefore establishes not only the optimal choice for our platform but also a set of design principles that can guide future therapeutic applications where maximizing editing efficiency is essential.

**Figure 3.**
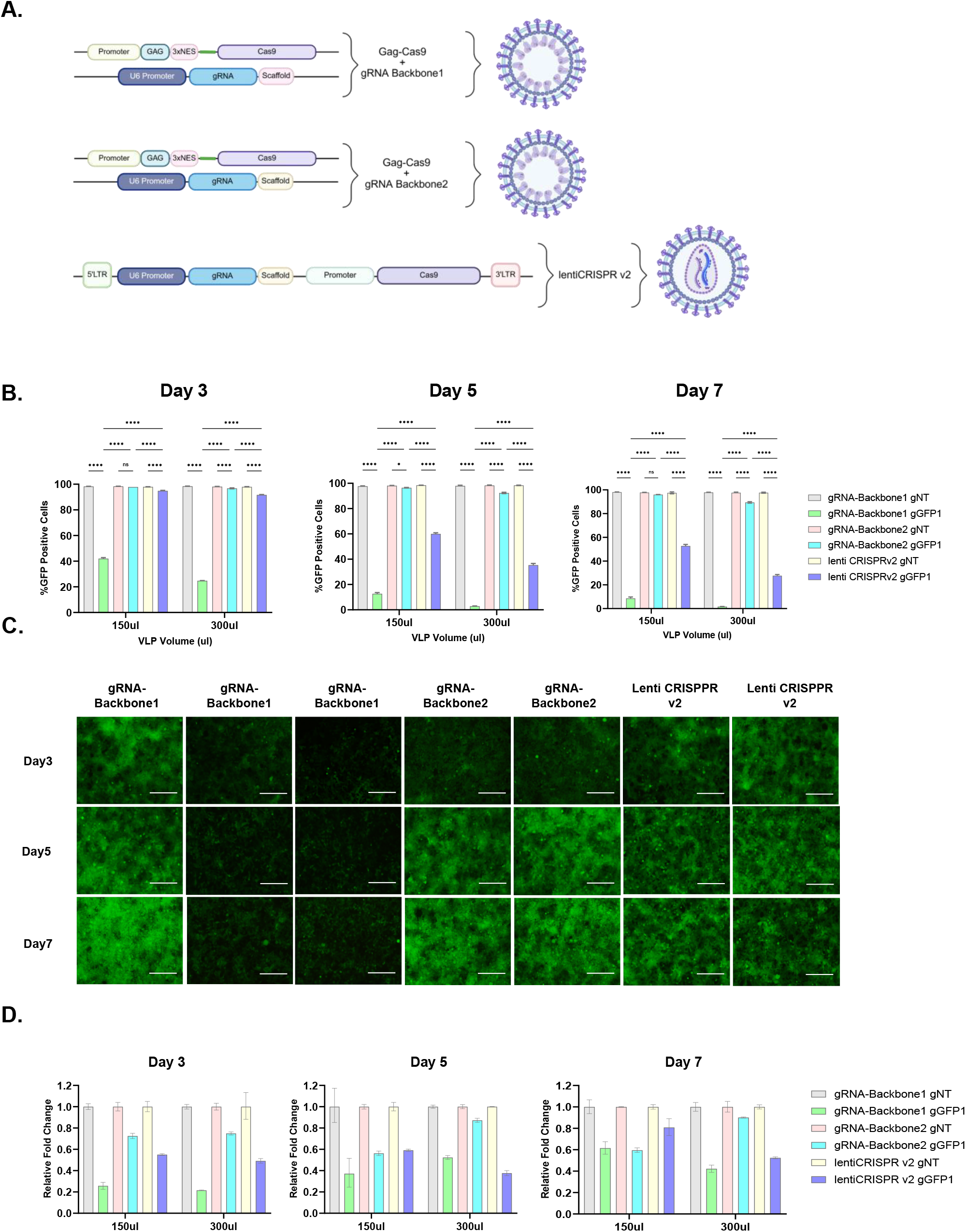
Comparison of gRNA delivery efficiency using different backbones and vectors. **A.** Schematic representation of gRNA backbone designs for VLP-mediated delivery. Two Gag–Cas9 fusion–based architectures were tested in combination with distinct gRNA backbones (Backbone 1 and Backbone 2), enabling RNP assembly and packaging into VLPs. For comparison, a conventional lentiviral construct (lentiCRISPR v2) was used, in which both Cas9 and gRNA are delivered as RNA. **B.** Flow cytometry analysis of GFP disruption following treatment with gRNA- Backbone1, gRNA-Backbone2, Lenti CRISPR v2-based systems at days 3, 5, and 7 using two different input volumes (150 µl and 300 µl). **C.** Representative fluorescence microscopy images showing GFP signal loss in reporter cells under the indicated conditions (scale bar, 200 µm). **D.** qPCR quantification of GFP transcript levels following delivery with gRNA-Backbone1, gRNA-BAckbone2, Lenti CRISPR v2-based constructs at days 3, 5, and 7, confirming the flow cytometry results. Statistical analysis was performed using two-way ANOVA. ^*^ p < 0.05, ^**^ p < 0.01, ^***^ p < 0.001, ^**^ p < 0.0001; ns: non-significant

### Cas9-RNP VLPs achieve potent and selective elimination of EGFRvIII-positive glioblastoma cells

To test whether targeted VLPs could translate efficient Cas9-RNP delivery into functional outcomes, we evaluated EGFRvIII-directed VLPs for their ability to induce selective cytotoxicity **(Fig. 4A).** VLPs coated with wild-type or mutant VSV-G served as non-specific controls, whereas scFv1-A and scFv1-V provided EGFRvIII-specific targeting modules. WT VSV-G–coated VLPs served as broad-entry controls, while mutant VSV-G provided reduced LDLR interaction; neither conferred EGFRvIII selectivity compared to scFv1-coated particles. We first confirmed genome editing efficiency using a gGFP1 reporter, where Cas9-RNP–loaded VLPs produced a five-fold increase in editing in EGFRvIII (+) cells compared to EGFRvIII (–) controls **(Fig. 4B).** This result demonstrated that receptor-targeted VLPs can deliver active nuclease cargo with both high efficiency and sharp discrimination between positive and negative populations.

**Figure 4.**
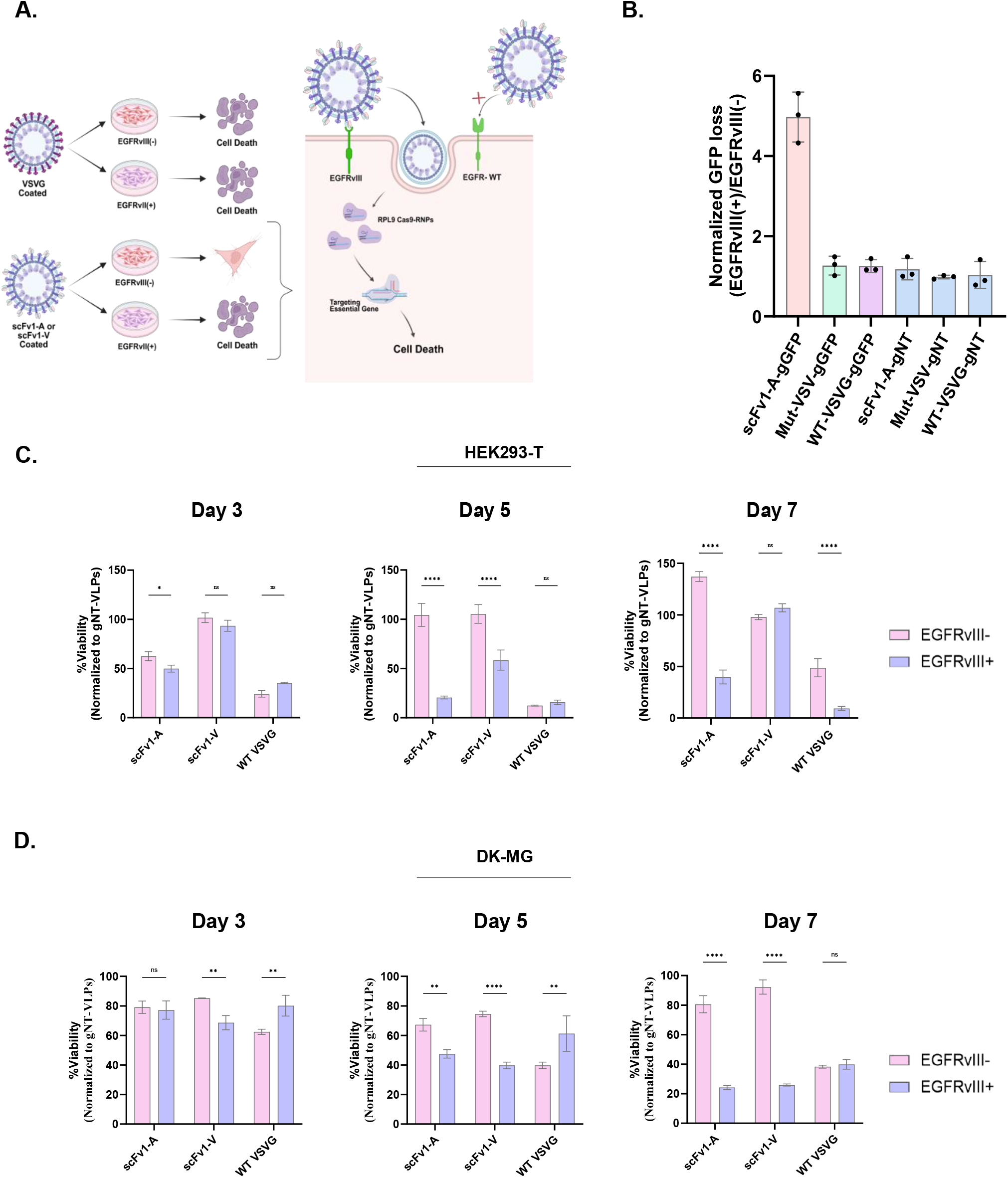
Targeted killing of EGFRvIII-positive cells using Cas9-RNP–loaded VLPs. **A.** Schematic representation of the expected outcomes upon treatment with VLPs coated with wild-type VSV-G, mutant VSV-G, or EGFRvIII-targeting scFvs (scFv1-A or scFv1-V). **B.** Editing efficiency of gGFP1-Cas9-RNP–loaded VLPs in EGFRvIII-positive cells, showing a five-fold increase compared with EGFRvIII-negative controls. Editing efficiency was normalized to GFP signal in EGFRvIII^−^ cells to account for background activity. **C.** Selective cell killing following delivery of Cas9-RNP targeting RPL9 in engineered HEK-293T EGFRvIII(+) versus EGFRvIII(–) cells. Viability was assessed at days 3, 5, and 7 by MTT assay. **D.** Selective cytotoxicity in DK-MG cells following treatment with Cas9-RNP–loaded VLPs coated with EGFRvIII-targeting scFvs, assessed at days 3, 5, and 7. Cell viability was normalized to NT gRNA–VLP treated cells. Statistical analysis was performed using two-way ANOVA. ^*^ p < 0.05, ^**^ p < 0.01, ^***^ p < 0.001, ^**^ p < 0.0001; ns: non-significant

Building on this, we next tested whether delivery of Cas9-RNPs directed against the essential gene, *RPL9,* could translate editing into cell killing. *RPL9* encodes a ribosomal protein of the 60S subunit that is indispensable for protein synthesis and has been consistently classified as a core essential gene in genome-wide CRISPR knockout screens[31]. In engineered HEK-293T reporter cells, EGFRvIII-targeting VLPs induced a striking loss of viability specifically in EGFRvIII (+) cells across all time points, while EGFRvIII (–) cells remained unaffected **(Fig. 4C).** The capacity to eradicate target-positive cells while sparing non-target cells is a defining benchmark for precision therapeutics, and our VLPs achieve this with remarkable clarity. Crucially, this selective cytotoxicity was reproduced in the endogenously EGFRvIII (+) DK-MG glioblastoma line **(Fig. 4D).** Demonstrating efficacy in this clinically relevant model provides strong validation that our engineered VLPs function not only in artificial reporter systems but also in patient-derived tumor cells, underscoring their translational promise. Together, these results establish that Cas9-RNP–loaded VLPs decorated with EGFRvIII-specific scFvs can achieve potent and highly selective elimination of glioblastoma cells bearing this tumor-specific receptor variant, while maintaining minimal effects on EGFRvIII-negative populations. This represents a critical advance toward programmable VLP-based therapeutics capable of discriminating tumor from normal tissue and delivering genome editors as precision cancer agents.

## DISCUSSION

In this work, we present a modular VLP platform capable of packaging Cas9-RNPs and selectively targeting glioblastoma cells that express EGFRvIII. Our findings demonstrate that rational engineering of VLP components—from entry modules to packaging strategies and gRNA backbones—can yield a system that combines efficient genome editing with selective tumor cell killing.

We first addressed the challenge of selective entry. By screening multiple scFvs and peptide ligands displayed via VSV- or ACE2-derived transmembrane domains, we identified scFv1 as the most effective targeting moiety. Importantly, the mode of display influenced not only efficiency but also specificity: VSV-TM provided the highest absolute entry levels, whereas ACE2-TM significantly reduced background uptake in EGFRvIII-negative cells, thereby enhancing discrimination. This modularity in entry design offers flexibility for adapting the system to different tumor-associated antigens beyond EGFRvIII [18, 32, 33]. Comparable scFv-based strategies have been used in EDVs to redirect viral particles toward T cells [10], and our data extend this approach by optimizing envelopes for a solid tumor–associated antigen.

We then optimized RNP packaging. Using a Gag–Cas9 fusion strategy, adapted from prior reports [8, 10, 13, 34], we achieved robust incorporation of Cas9 protein into VLPs. Systematic titration of plasmid stoichiometry revealed that efficient genome editing was obtained across a wide range of ratios, but maximal and reproducible outcomes were achieved at balanced conditions (Gag-Cas9:pUMVC ratios; 1:1, 1:2, 2:3). Notably, excessive Cas9 input increased incorporation of protein but was accompanied by reduced VSV-G levels, indicating a stoichiometric imbalance that compromises particle quality. This highlights the importance of maintaining equilibrium between structural and cargo components during VLP production. Such systematic optimization is rarely performed, yet it is critical to establish reproducible parameters that can bridge the gap between proof-of-concept and translational application.

A critical component of our workflow was the use of a homozygous GFP knock-in line at the hROSA26 locus. This system provided a quantitative and scalable readout of editing efficiency and allowed us to cross-validate results obtained by flow cytometry and qPCR. Reporter systems based on safe-harbor integration have previously been shown to offer stable and reproducible editing readouts [35], and our results confirm that such models are well suited for benchmarking VLP delivery.

We further investigated the influence of gRNA backbones. Comparison between gRNA-backbone1 and gRNA- backbone2 constructs revealed that backbone1 conferred consistently higher editing efficiency, both in engineered cells and in disease-relevant DK-MG glioblastoma cells. Moreover, under the tested input volumes, VLP-mediated delivery outperformed the conventional lentiviral vector lentiCRISPR v2, which packages Cas9 and gRNA as RNA. This observation is consistent with earlier findings that scaffold structure significantly impacts gRNA stability and editing outcomes [36]. Although absolute particle numbers were not normalized in this comparison, the consistent superiority of backbone1 VLPs across multiple readouts and time points suggests that this backbone is particularly well-suited for RNP delivery applications. Systematically benchmarking gRNA backbones in the context of VLP-mediated RNP delivery is rarely undertaken, yet it is essential to define design principles that ensure reproducibility and translational relevance beyond proof-of-concept studies.

Most critically, we demonstrated that EGFRvIII-targeted VLPs can translate efficient delivery into functional outcomes. When loaded with Cas9-RNPs targeting RPL9, scFv1-coated VLPs induced potent and selective killing of EGFRvIII (+) cells, while sparing EGFRvIII (-) controls. This effect was observed both in engineered HEK-293T cells and in the endogenously EGFRvIII (+) glioblastoma line DK-MG, underscoring the potential clinical relevance of our approach [37]. RPL9 encodes a ribosomal protein of the 60S subunit, indispensable for protein synthesis, and ribosomal proteins are consistently classified as core essential genes in genome-wide dependency maps [31]. Thus, the ability to eliminate EGFRvIII-positive cells by targeting such an essential gene provides a stringent demonstration of selective cytotoxicity. By linking a well-defined essential gene to receptor-targeted delivery, our study goes beyond showing editing efficiency and directly proves that engineered VLPs can translate genome modification into functional and therapeutically meaningful cell elimination.

Despite these promising results, several limitations should be acknowledged. While we demonstrated efficient editing and selective cytotoxicity *in vitro, in vivo* validation remains essential to assess biodistribution, blood– brain barrier penetration, tumor retention, and safety. Strategies such as convection-enhanced delivery or intratumoral injections may be required to achieve effective dosing in brain tumors [38-41]. Immunogenicity also remains a consideration: although VLPs are non-replicating and Cas9 delivery is transient, both the structural proteins and the bacterial Cas9 nuclease could elicit immune responses *in vivo* [42-44]. Strategies such as employing orthogonal nucleases or engineering Cas9 variants with reduced immunogenicity are being actively investigated [45]. In addition, alternative transmembrane domains in place of canonical VSV-G may be explored to maintain fusogenic activity while potentially reducing immune recognition [46]. Finally, optimization of large-scale production and purification will be critical for eventual translational applications [47].

Looking ahead, our findings establish a framework for the development of targeted VLP therapeutics. The modularity of the system means that other tumor-specific scFvs or peptide ligands could be readily substituted to expand the range of indications. Moreover, while this study focused on genome editing as a mechanism of killing, the same platform could be adapted to deliver alternative protein cargos such as pro-apoptotic enzymes or immune-modulating factors [48, 49]. Beyond oncology, targeted RNP-VLPs could be employed for transient, tissue-specific genome editing in regenerative medicine, metabolic disorders, or immunotherapy [34, 49, 50].

In conclusion, we demonstrate that Cas9-RNP–loaded VLPs decorated with EGFRvIII-specific scFvs enable potent and selective elimination of glioblastoma cells *in vitro*. This work advances the concept of programmable VLPs from a delivery tool toward a therapeutic modality and provides a framework for future preclinical evaluation and potential therapeutic translation of targeted protein- and RNP-based therapies.

## ACKNOWLEDGMENTS

Financial support was obtained from Health Institutes of Türkiye (TUSEB)(33335 Grant). The authors acknowledge the financial support and use of the services and facilities of the Koç University Research Center for Translational Medicine (KUTTAM). C.B-O. is supported by a TUBITAK-BIDEB 2211 scholarship for PhD studies.

